# TRAILBLAZER: generative multicellular perturbation model of biology

**DOI:** 10.64898/2026.03.14.711710

**Authors:** Julian Neñer, Padmalochini Selvamani, Smitha Srinivasachar Badarinarayan, Naveen Chandramohan, Adrian Grzybowski

**Author notes:** Corresponding author: A.G.

## Abstract

Single-cell foundation models are reshaping biology by learning transferable representations of cellular state from millions of profiles. These models support annotation, denoising, cross-modal mapping and, increasingly, prediction of responses to genetic or pharmacological perturbations. Despite this progress, most approaches treat cells as independent observations and ignore the multicellular context that governs tissue behavior. Models trained on aggregated datasets often fail to generalize to new donors, laboratories or interventions, in part because their latent spaces lack structure for composition and extrapolation. As a result, strong reconstruction performance does not guarantee accurate forecasting of system-level responses.

The general problem addressed here is how to construct a scalable model that predicts multicellular, patient-level responses to interventions while preserving single-cell resolution and enabling generalization beyond observed conditions.

Here we show that TRAILBLAZER, a multicellular transformer encoder coupled to an explicitly shaped hyperspherical latent space and a count-aware generative decoder, enables accurate zero-shot prediction of perturbation responses and ranking of candidate immunomodulators at patient resolution.

In contrast to prior single-cell or pseudo-bulk approaches, TRAILBLAZER models tissues as coordinated systems using latent tokens that summarize and redistribute global context while maintaining near-linear scaling with group size. By organizing latent geometry around shared healthy references and calibrated mechanistic directions, the model renders vector arithmetic biologically meaningful and supports extrapolation to unseen agents. Together, these results establish a practical framework for mechanism-aware simulation of multicellular responses and suggest a path toward predictive foundation models for therapeutic discovery.

## INTRODUCTION

Single-cell foundation models have transformed how we represent cellular state and reuse knowledge across studies. Pretrained on millions of profiles, these models learn transferable embeddings that support core tasks such as automated cell-type annotation^1^, label transfer across cohorts and chemistries^2^, denoising^3^, imputation^4^, cross-mapping data modalities^5^, and increasingly, forecasting the effects of perturbations, such as pharmaceutical interventions, environmental changes, genetic edits. Methodologically, they span probabilistic count models (e.g., VAE-based)^6^, large transformer encoders^7^, and generative frameworks adapted from vision^8^ and language models^9^. Together they promise a common computational substrate for discovery, enabling downstream analysis to start from rich priors.

Despite this progress, important limitations remain. Most single-cell foundation models treat cells as independent and identically distributed observations, discarding the multicellular context that, in vivo, shapes system-level outcomes^10^. Models trained on aggregated datasets often fail to generalize under unseen conditions, such as new donors, laboratories, chemistries, tissues, or interventions, due to training distributions being narrow and the learned latent representations lacking the structure required for composition (the ability to combine known elements in novel ways) and extrapolation (the ability to predict beyond the range of observed data).

Multicellular models aim to address these gaps by modeling tissues as coordinated dynamical systems rather than “bags of cells”^11^. In real tissues, intercellular signaling and feedback give rise to stable attractors that constrain and guide individual cells^12–14^. Synergistic effects across cell types are commonplace^15,16^; conversely, any one cell’s state may be noisy or weakly informative.^17–20^ Encoding groups of cells and allowing information to flow among them therefore captures macroscopic constraints that stabilize phenotypes and shape responses to interventions.

Recent multicellular encoders begin to address this need by operating on unordered sets of cells. Previously, deep sets-style aggregation^21^ has been employed to embed each cell independently and pool them together with a permutation-invariant operator (linear *O(N)*). Approaches in this family including PaSCient^22^ and CellFlow^23^ demonstrate that set-level reasoning improves patient-level characterization, yet pseudo-bulk pooling can be sensitive to group composition and tends to discard per-cell detail needed for reconstruction and simulation. More recently, STACK^24^ moves beyond pooling by applying tabular attention across cells in a shared context window, allowing each cell’s representation to be informed by its co-profiled neighbors. However, scaling attention beyond thousands of cells typical of patient-level samples remains a practical challenge. Explicit all-pair cell-cell attention/message passing would, in principle, be capable of capturing interactions but is *O(N²)* and therefore computationally costly and prone to overfitting.

In contrast, we use permutation-invariant, transformer-based encoders where learned latent (inducing) tokens summarize global cellular context and feed it back to recontextualize individual elements, providing an explicit information bottleneck^25^. This design preserves two-hop global cellular context while keeping compute and memory effectively near-linear in group size, enabling stable training and inference on large cell sets and making multicellular modeling practical for routine perturbation prediction.

Orthogonal to the encoder, we argue that the geometry of the latent space must be deliberately organized. In computer vision^26^, speech^27^, and language^28^, metric learning has shown that hyperspherical embeddings with angular objectives and contrastive structure make directions semantically meaningful and composable^29^. We propose to use the same principle for models of biology. When such geometry is enforced, vector arithmetic becomes valid, one can add or subtract interventions, project a desired change onto a library of mechanisms, and generalize to unseen agents in zero- or few-shot settings, while retaining a faithful generative model of per-cell counts.

Here, we introduce TRAILBLAZER, which unites a scalable multicellular encoder with an explicitly shaped latent space and a count-aware decoder to simulate how patients’ cell populations respond to interventions. Upstream, a harmonization module removes batch artifacts; a separate mechanism-segmentation network, trained before TRAILBLAZER, provides a frozen library of mechanisms directions that anchors latent shaping; and a non-trivial dataloader constructs donor-matched, cell-type-balanced groups at scale to prevent composition shortcuts. We show that TRAILBLAZER accurately predicts multicellular molecular profiles and phenotypes in zero-shot settings, rediscovers and ranks immunomodulators effective in treating cancer, and that it scales linearly with group size while improving reconstruction as more cells are considered. Across benchmarks, including comparisons with recent single-cell and multicellular baselines, TRAILBLAZER preserves full-distribution properties, generalizes beyond training conditions, and produces ranked treatment suggestions at patient resolution.

## RESULTS

### Overview of TRAILBLAZER

TRAILBLAZER is a generative model that enables a range of downstream biological and translational analyses by modeling multicellular responses and phenotypes to perturbations across immune-related disease settings (Fig. 1a). The model facilitates phenotypic drug discovery by learning treatment-induced state transitions across interacting cell populations. This approach enables treatment prioritization based on phenotypic multicellular outcomes, without reliance on prior specification of molecular targets. Furthermore, TRAILBLAZER helps predict and simulate virtual patient digital twins’ responses to treatments that are useful for clinical trial simulation, supporting rational trial design, and cohort selection. Lastly, our framework enables more detailed analyses, including cell-type and feature importance, to determine which cellular populations and biological programs most strongly contribute to the observed phenotypic differences (Fig. 1b).

**Fig. 1.**
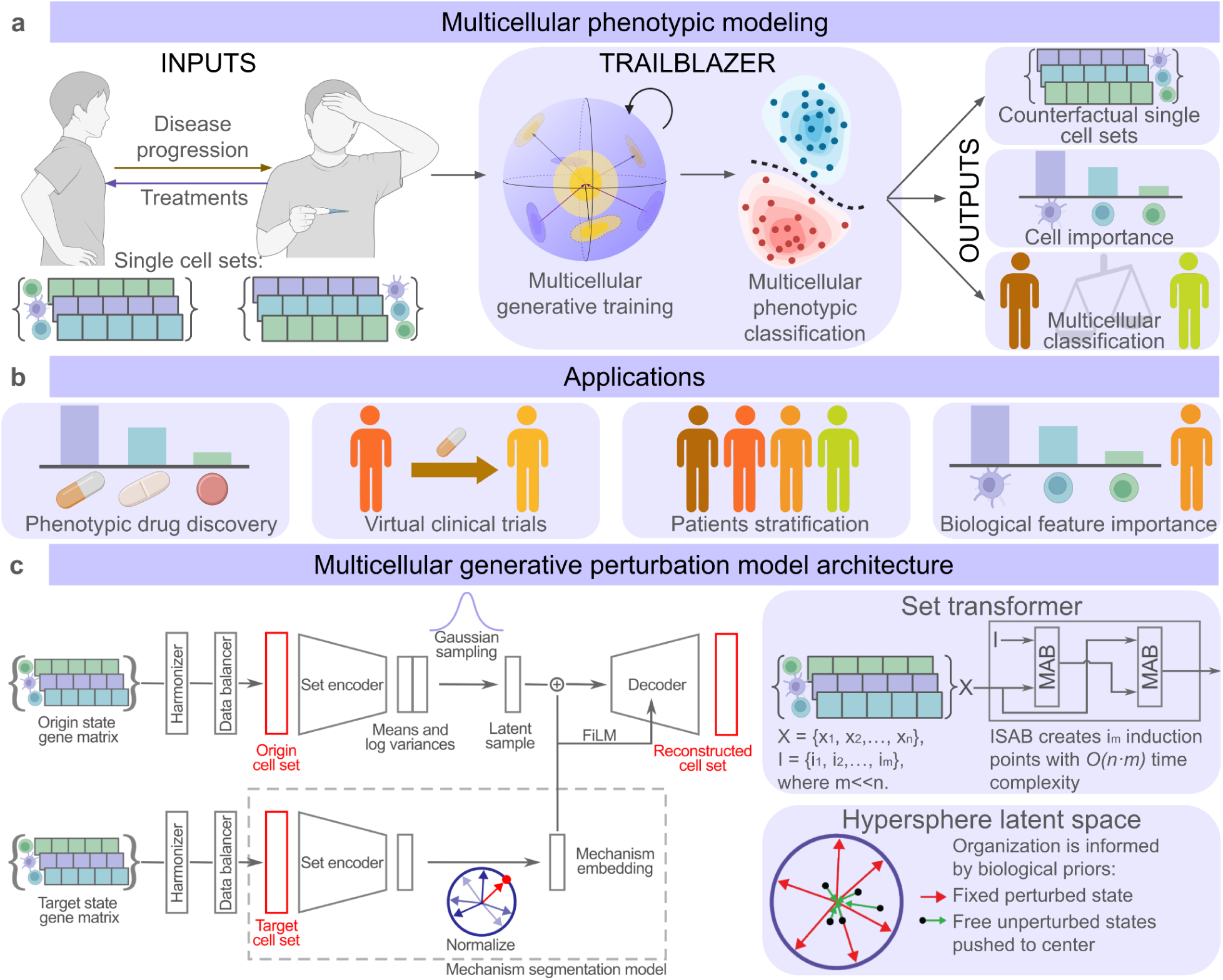
TRAILBLAZER: Multicellular architecture for foundational models of biology. **a,** In multicellular organisms, sets of cells are working together to produce phenotypic presentations such as treatment response and disease progression. TRAILBLAZER operates on sets of cells to build uniform and traversable hypersphere latent representation of multicellular disease and treatment processes; it is coupled with multicellular classifiers to help with navigating the phenotypic space. **b,** Uniform disease-treatment phenotypic perturbation space can be applied to discover treatments by presenting initial and target patient phenotypes, or it can be used to identify patients phenotypes in response to treatments in virtual clinical trials, lastly multicellular architecture offers unbiased view of each cell importance to the process. **c,** TRAILBLAZER uses Induced Set Attention Blocks (ISAB)^25^ to efficiently capture relationships of large sets of cells’ biological variables such as scRNA-seq, and utilizes latent shaping with biological priors to improve generalizability by organizing latent space on a plan of hypersphere with healthy states in the center.

To model population level effects, TRAILBLAZER employs a permutation-invariant multicellular encoder in which learned latent (inducing) tokens aggregate global cellular context and redistribute it back to individual cells. Furthermore, TRAILBLAZER deliberately shapes the geometry of its latent space to make biological interventions composable and interpretable. Presented here architecture enables zero- and few-shot generalization to unseen treatments while maintaining a faithful generative model of per-cell molecular counts (Fig. 1c).

### Loading and preparing single-cell data

All inputs are first mapped into a common gene index (Supplemental data 1) (comprising highly variable genes plus key markers) and, if needed, harmonized so that batch effects are removed upstream while preserving biological variation. We assemble training examples as donor-matched pairs of cell sets: one set of unperturbed cells (e.g., vehicle or healthy) and one set of perturbed cells (treatment or disease). Matched by donor to cancel idiosyncratic baseline differences that would otherwise dominate the signal. No order is imposed on the cells in the set. Crucially, each pair is balanced by cell type so the model is forced to learn within-cell-type transcriptomic shifts due to the perturbation rather than trivially exploiting changes in cell-type composition between conditions. To determine an effect of cell balancing on cell reconstruction accuracy, we have trained two networks on PARSE PBMCs perturbation dataset^30^. The networks were trained using sets of 500 cells with the task of reconstructing cells; using 9 donors for training and 3 donors for validation (unseen). The post-treatment transcriptomic profiles were reconstructed from input cells latents, and average perturbation embeddings produced with the mechanism segmentation network. It was observed that cell balancing improves cell reconstruction of all cell types in the population (Ext. Fig. 1a,b).

### Encoding sets of cells

Each cell set is embedded by a permutation-invariant transformer encoder that allows information to flow among all cells while keeping compute and memory close to linear in set size. Communication is routed through a small number of learned relay (inducing) tokens; in the first pass, relays attend over all cells to absorb global context; in the second, cells attend back to the relays to receive context-aware updates (Fig. 2a). This two-hop design preserves long-range interactions without incurring quadratic costs (*O(n·m)* with m relays, m ≪ n), enabling thousands cells per forward pass on standard GPUs. The encoder outputs a latent vector for every cell together with a compact context token that summarizes sample-level information used later for conditioning and latent arithmetic. Pre-norm residual blocks, masking, and layer normalization stabilize training across variable set sizes. Empirical latency and memory scaling with cell set size are reported in the (Ext. Fig. 2a,b). The essential property of the set encoder is that it preserves intercellular dependencies, inaccessible to single-cell models, while remaining computationally tractable for clinically sized samples.

**Fig. 2.**
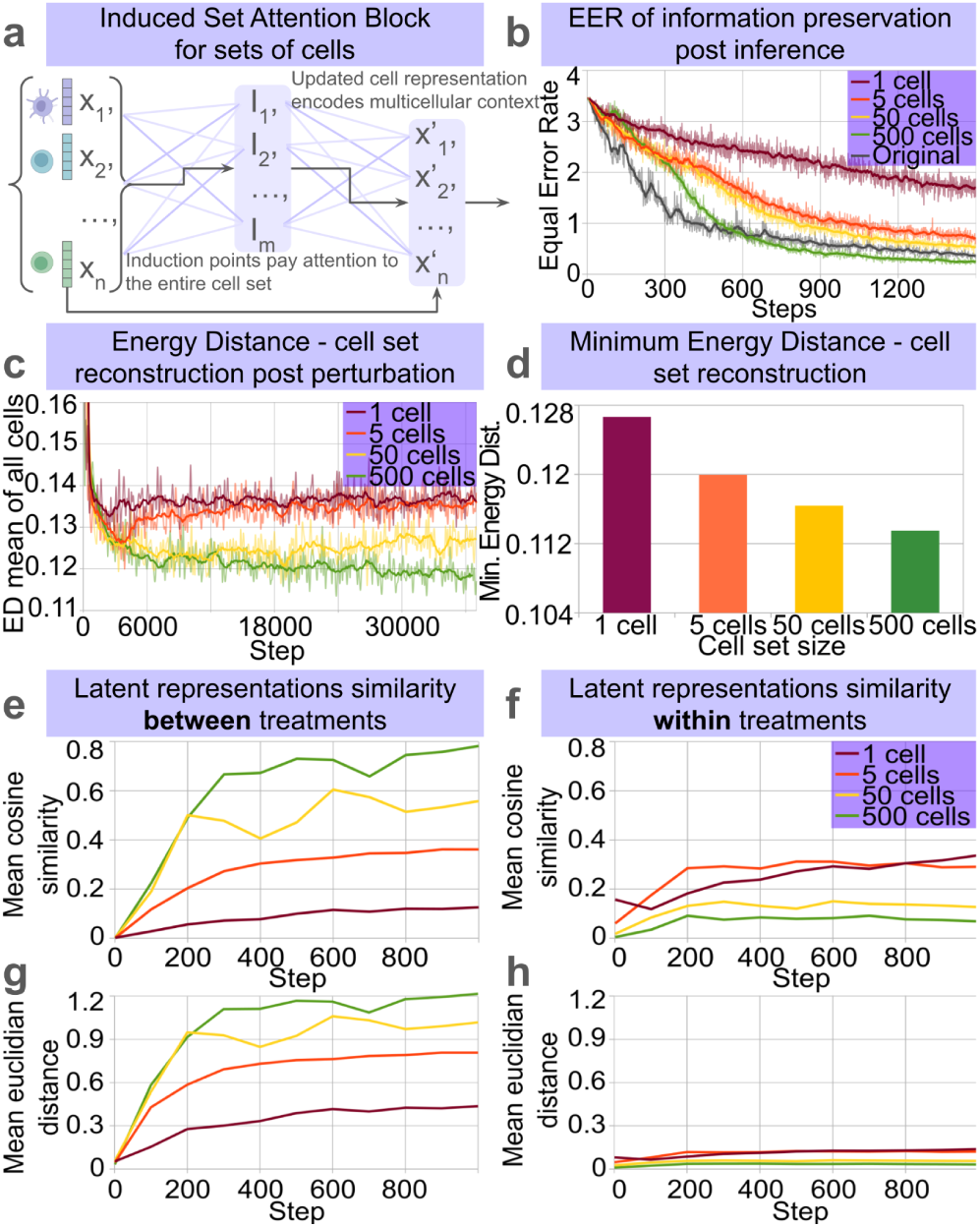
Set transformers as attention-based permutation-invariant encoders for cell sets. **a**, Induced Set Attention based encoders capture all-to-all cell interactions by leveraging induction point attention, while reducing time complexity from *O(n^2^)* to *O(nm).* **b**, Equal error rate (lower better) for correct treatment classification based on fixed cell set size (50 cells) for cell sets scRNA-seq inferred by networks trained on groups of 1, 5, 50 and 500 cells, “original” set indicates ground truth cell set. **c**, Energy distance of genes, mean of 25 cells per cell type, for reconstruction of scRNA-seq of unseen donors (N=3), calculated every 100 steps of the training on sets of 1, 5, 50 and 500 cells, for 90 treatments and 9 donors. In all runs, the network sees exactly 1000 cells in each forward pass, independent of cell set size. Latent shaping was not applied. **d**, same as c, but the minimal energy distance for each cell set size. **e**, the network was trained on the task of clustering sets of cells by treatment, trained on sets of 1, 5, 50 and 500 cells, for each run batches have the same amount of states to cluster per forward pass. The cosine similarity was measured for cell samples between treatments (higher better), or **f**, within treatment (lower better). **g**, Same as (f) but mean euclidean distance for cell samples between treatments (higher better), or **h**, within treatment (lower better). All measurements use the PARSE PBMCs dataset.

To verify that TRAILBLAZER preserves intervention specific information in reconstructed cells, we leverage the mechanism segmentation network (described in the following section) as a downstream probe. Because this network is trained to separate and cluster cells according to their intervention, its performance on synthetic data directly reflects how faithfully the reconstructed latent space retains biological signal. Concretely, we trained and inferred TRAILBLAZER on the PARSE PBMCs dataset^30^ using sets of 1, 5, 50, and 500 cells, then trained the mechanism segmentation network independently on each resulting synthetic dataset, using sets of 50 cells in all cases. As shown in Fig. 2b, models conditioned on larger cell sets yield systematically lower Equal Error Rate (EER^31^), indicating that richer cellular context leads to reconstructions that better preserve the structure needed to distinguish interventions.

Reconstruction quality as measured by mean energy distance further corroborates this trend, decreasing consistently as context set size grows (Fig. 2c,d; Ext. Fig. 2c), suggesting that larger groups allow TRAILBLAZER to extract richer contextual signals and generate more faithful reconstructions.

### Mechanism segmentation network

Prior to training TRAILBLAZER we trained a separate “mechanism segmentation” model that supplies the intervention directions used to anchor the latent space. This model takes sets of cells from a single donor under a single intervention and embeds each cell set with the permutation-invariant set encoder described above. It is trained with a similarity-matrix objective on a hypersphere (GE2E-style)^31^, pulling together embeddings of the same intervention across donors and pushing apart different interventions; embeddings are constrained to unit norm.

Consistent with abovementioned improvements associated with training on increasing context cell set sizes, an analysis of inter- and intra-group distances in the latent space reveals an improved perturbation clustering; as cell set size increases, both euclidean and cosine distances between cells from different treatments grow (Fig. 2e, 2g), while distances between cells from the same treatment shrink (Fig. 2f, 2h).

After training, we form a frozen dictionary by averaging per-donor embeddings for each intervention and re-normalizing, as well as fitting a von Mises–Fisher distribution to capture dispersion across donors. Conceptually, this produces one unit vector per mechanism together with a concentration parameter, i.e. a library of calibrated directions in which nearby vectors correspond to biologically related actions. This library is not updated during TRAILBLAZER training; it serves as a stable coordinate system for latent shaping and for downstream ranking.

### Latent shaping

Latent shaping organizes TRAILBLAZER’s representation so that geometry reflects biological mechanisms. Given per-cell latents x for a control group and y for a perturbed group from the same donor, and a mechanism vector g from the frozen mechanism segmentation library, we perform latent arithmetic in both directions: x is shifted toward x+g to simulate the perturbed state, and y toward y−g to simulate removal.

Practically, training proceeds in mini-batches of donor-matched pairs of sets, one control and one perturbed, with cell type balanced sampling and a fixed product of batch size and group size so that gains from larger sets reflect genuine multicellular reasoning. For each pair we form four decode paths that expose both reconstruction and simulation behavior in the same update: control→control, perturbed→perturbed, control→(apply g)→perturbed, and perturbed→(remove g)→control. We encourage the displacement between control and perturbed embeddings to align with g via a cosine objective, while gently compacting healthy controls toward a shared reference near the origin and, when needed, normalizing perturbation magnitudes so directions have comparable length. In simple terms, healthy, unperturbed states are pushed to the center of the hypersphere and sick and perturbed states are put on the surface of the hypersphere (Fig. 3a).

**Fig. 3.**
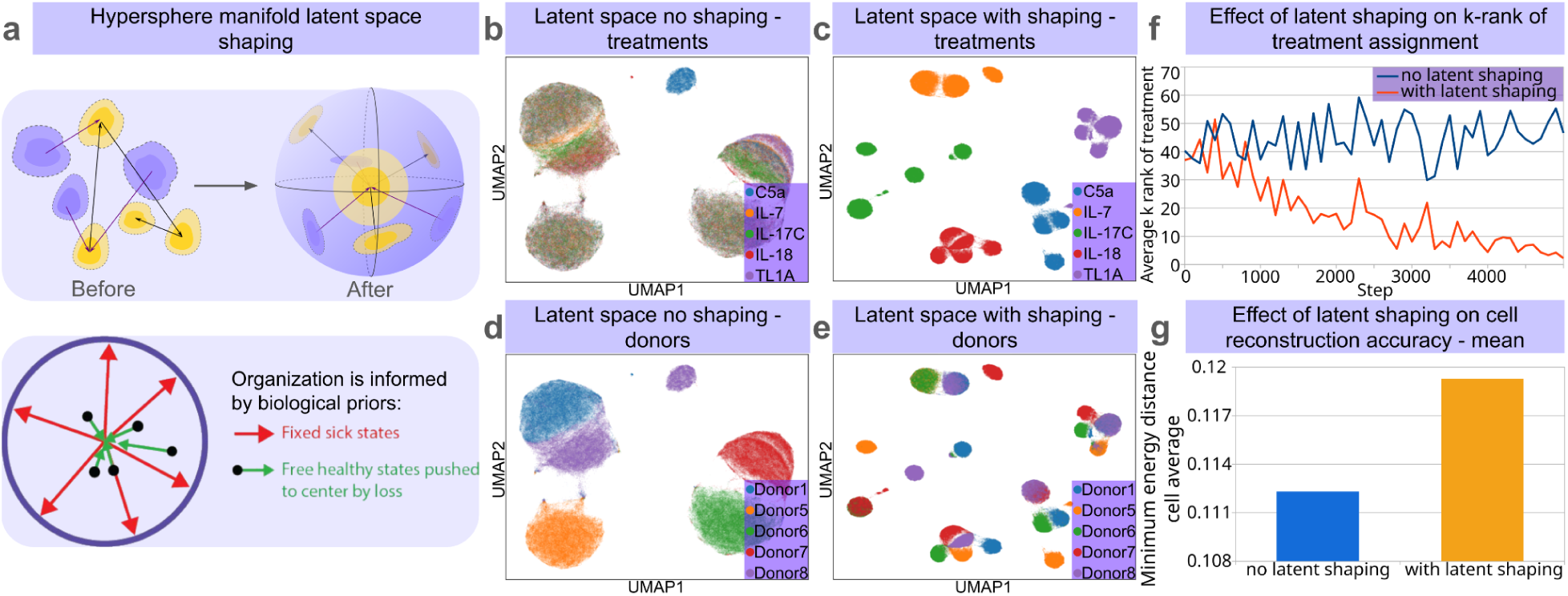
Application of latent space shaping using biological priors. **a**, To improve generalizability of TRAILBLAZER, the latent space of the network is shaped using a “Mechanism segmentation network” trained on a hypersphere manifold, and where healthy, unperturbed states are pushed towards the center and sick and perturbed states are put on the surface of the hypersphere. **b**, UMAP projection of the latent space for PARSE dataset’s 5 random treatments without latent shaping, and **c**, with latent shaping. **d**, same as b but view of 5 random donors without latent shaping, and **e,** with latent shaping. **f**, Average k-rank of treatment rediscovery for unseen donors based on cosine similarity of latent embeddings (lower better). **g**, Minimum energy distance for network without and with latent shaping calculated for each gene, averaged across 25 cells per cell type, for reconstruction of scRNA-seq of unseen donors (N=3), trained on sets of 500 cells. All measurements use the PARSE PBMCs scRNA-seq dataset.

Directly optimizing the full objective by combining reconstruction losses and latent metric regularization from the start was found to lead to suboptimal convergence. For that reason, we adopt a staged training that separates the hard parts of the problem. In the first stage we train reconstruction and apply only the radial term for controls, allowing the encoder–decoder to settle while anchoring the healthy reference. In the second stage we introduce the angular (cosine) alignment so that perturbation directions behave consistently across donors and tissues; empirically, this is the point at which latent representations pivot from donor-dominated to perturbation-dominated structure, and the rediscovery rank improves sharply. In the third stage we add the optional norm constraint on displacement so vector composition (e.g., g1+g2) becomes stable across batches. Throughout training we monitor a cosine-matrix distance between TRAILBLAZER’s latent means (per donor × perturbation) and the segmentation reference computed on the same samples; when this distance reaches a minimum we freeze the encoder and continue decoder-only fine-tuning. Freezing encoder at geometric optimum preserves the best latent geometry while allowing the count head to improve per-gene fit without dragging the space away from its useful alignment.

### Latent shaping enables discovery and generalization

Latent shaping reorganizes the representation so that geometry reflects biology rather than donor identity. To visualize this effect, we inferred latents for 5 randomly selected donors and 5 treatments before and after training with the shaping objectives, and projected resulting latents with UMAP. In the unshaped space, embeddings cluster primarily by donor with little visible structure by treatment, indicating that inter-donor idiosyncrasies dominate the geometry (Fig. 3b,d). After shaping, the organization flips with treatment-specific clusters emerging (Fig. 3c,e). This qualitative shift is the intended effect of aligning angular directions with mechanisms.

We quantified the impact of latent shaping on treatment re-discovery by cosine-projecting the mean latent of each donor-treatment pair onto the frozen mechanism library and averaging the rank of the correct treatment across pairs. Without shaping, the rediscovery rank is approximately random. With shaping, the average rank drops steadily during training (Fig. 3f), indicating that mechanism directions have become usable for projection and ranking. This improvement is robust across group sizes and donors and co-occurs with the qualitative transition in latent representation from donor-centric to treatment-centric, with only a modest trade-off in energy distance of the cell’s transcriptome reconstruction (Fig. 3g, Ext. Fig. 3). This pattern is consistent across donors and group sizes and reflects the intended reallocation of capacity from purely local reconstruction to a geometry that supports projection, composition, and generalization.

Shaping also enables generalization to unseen interventions. In a zero-/few-shot benchmark on the PARSE PBMCs^30^ dataset, we held out IL-15 entirely during training and matched state-of-the-art phenotypic perturbation models, including CellFlow^23^, STACK^24^, and LPM^32^, on the same 5634 gene space in a subset of 1M cells. For the zero-shot predictions, TRAILBLAZER achieves substantially higher reconstruction accuracy than all competing architectures when predicting IL-15-treated cells from the same donors’ controls by adding the IL-15 mechanism vector and decoding (Fig. 4). The magnitude of this advantage demonstrates the importance of deliberate latent shaping. Interestingly, adding a small number of IL-15 donors during training (few-shot) yields little additional gain, suggesting that once the latent geometry is shaped, the model can compose the unseen direction without direct supervision on that treatment.

**Fig. 4.**
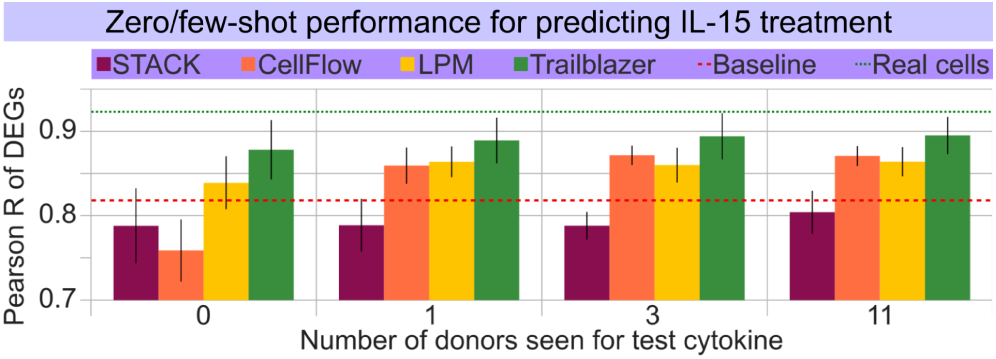
TRAILBLAZER’s architecture exhibits superior performance for both zero-shot and few-shot treatment outcome prediction. Post-treatment reconstruction Pearson R score for top 100 differentially expressed genes in a task of predicting post-treatment scRNA-seq state of the withheld donor for networks trained on 0, 1, 3 and 11 donor examples of the test treatment. Baseline control indicates no change from pretreatment state, real cells denote ground truth, sampled in the same fashion as predictions. Networks are trained on a common dataset of ex vivo 90 treatments of PBMCs from 12 donors. Measurements are means of 100 samples and their standard deviation. Comparison of STACK, LPM, CellFlow, and TRAILBLAZER models performance for IL-15 treatment prediction. All measurements use the PARSE PBMCs scRNA-seq dataset.

We subsequently repeated the experiment for IL-10 (Ext. Fig. 4a) and IFN-γ (Ext. Fig. 4b) and observed similar trends. CellFlow’s zero-shot model remains below the identity baseline in our setup and in accordance with previous findings^23^, indicating that its embedding of unseen perturbations without explicit supervision does not transfer as effectively. STACK also falls particularly behind in this low-data regime, as its masked reconstruction pretraining objective requires exposure to significantly more cells to learn generalisable representations. In contrast, LPM achieves better than baseline performance for zero-shot and few-shot reconstructions, but remains below that of TRAILBLAZER.

### Patient stratification, virtual trials, and discovery

We evaluated TRAILBLAZER as a virtual trial engine in patients with non-metastatic, treatment-naive primary invasive carcinoma of the breast treated with one dose of pembrolizumab (Keytruda or anti-PD-1) approximately 9 ± 2 days before surgery^33,34^. For this task we have coupled TRAILBLAZER’s counterfactual single-cell transcriptomic simulations with a multicellular classifier using the same multicellular encoder as the TRAILBLAZER (Fig. 5a). The classifier operates on sets of cells (100–500 per evaluation), aggregates information across the population, and outputs a patient-level probability of responsiveness defined as TCR clonotype expansion as in Bassez et al., 2021; trained on post-treatment states and evaluated on held-out donors, it achieves strong discrimination in the absence of TCR-seq data (ROC AUC ≈ 0.93; Fig. 5b). We then applied TRAILBLAZER to naive, pre-treatment samples from unseen patients by providing patients pre-treatment scRNA-seq along the α-PD-1 mechanism vector and decoding the counterfactual post-treatment cells. Running the multicellular classifier on these simulated cell sets reproduced treatment responsiveness of the unseen patients as compared to clinical ground truth (compare Fig. 5c and 5d). As a negative control, we performed the same procedure with IL-10 and observed the expected broad suppression of predicted response across patients (Fig. 5e), consistent with IL-10’s tolerogenic effects^35–37^.

**Fig. 5.**
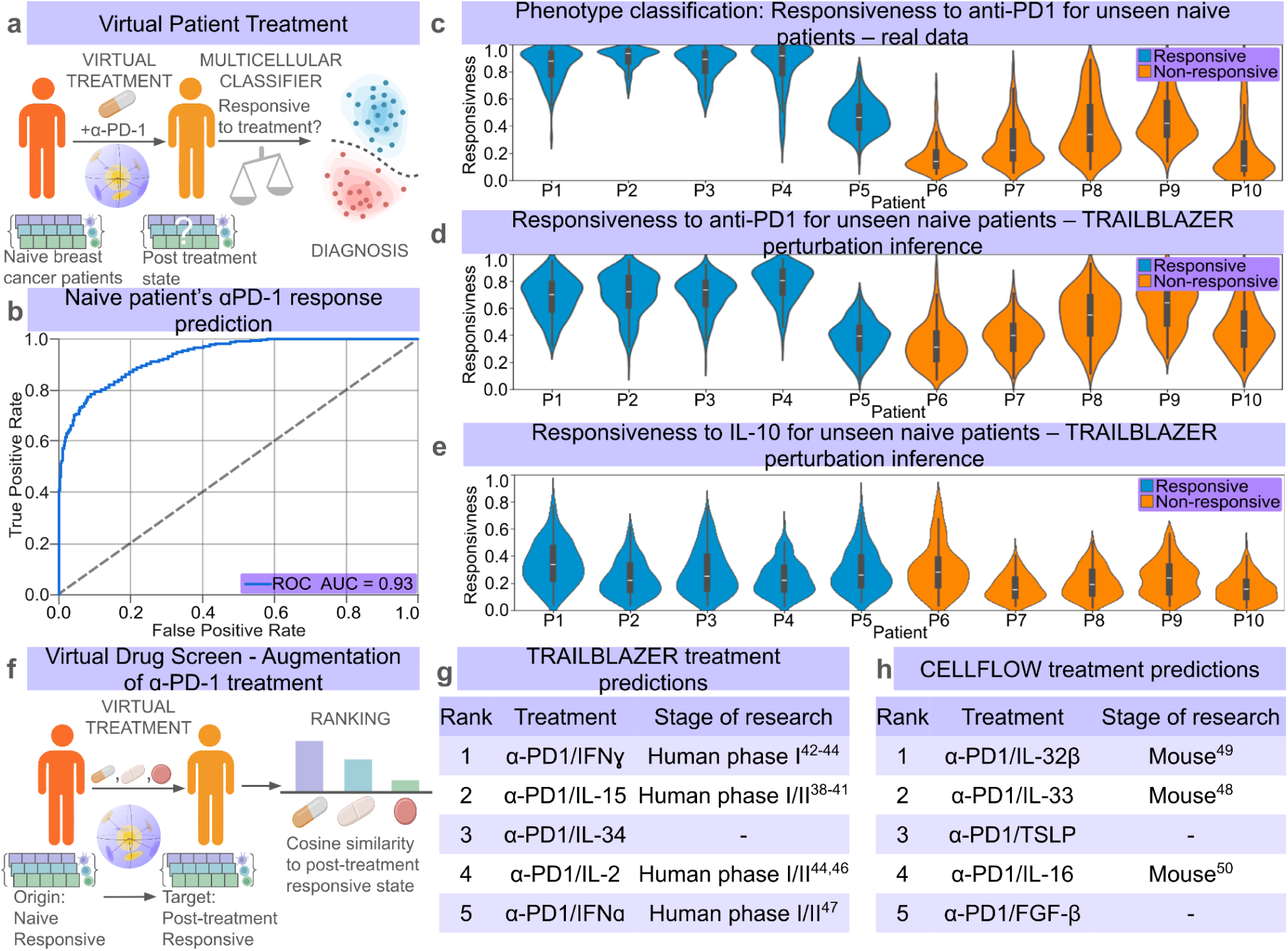
TRAILBLAZER stratifies individual patients’ response to α-PD-1 treatment. **a**, Graphical abstract for virtual patient treatment, scRNA-seq data for an unseen group of patients is passed to the perturbation model to generate counterfactual dataset subsequently passed to multicellular classifier to measure efficacy of the treatment, pertinent to figures (b-e) for virtual treatment of breast cancer patients with anti-PD-1. **b**, ROC curves for predicting the response of naive breast cancer patients to α-PD-1 treatment. **c**, We trained a multicellular respondent/non-respondent classifier on a single-cell α-PD-1 dataset, labels reflect eventual clinical response to the anti-PD-1 treatment. For each patient, we predicted response probabilities for samples of 100 cells, repeated sampling 100 times, and showing underlying distribution as a violin plot. **d**, We have used TRAILBLAZER to apply α-PD-1 treatment in silico and repeated classification as in (c). **e**, As in (d) but after IL-10 treatment. **f**, Graphical abstract for virtual drug screening, initial and target multicellular states are used to calculate a phenotype vector compared to mechanism vectors with cosine similarity to rank treatments most likely moving in direction of desired health outcome. **g**, TRAILBLAZER’s top five zero-shot predictions for treatments augmenting ɑ-PD-1 response, derived from scRNA-seq biopsies of breast cancer patients treated with ɑ-PD-1. Success was assessed by literature review corroborating treatment efficacy. Predictions are shown for unseen patients. **h**, Same ranking task as in (g), performed with the CELLFLOW neural network on the identical data foundation, harmonized with TRAILBLAZER’s harmonizer module.

Beyond single-agent simulation, TRAILBLAZER supports patient-specific ranking of α-PD-1 combinations. For each naive patient, we estimate the required delta that steers the observed pre-treatment state toward a post-treatment α-PD-1 responder target state in latent space, then project that delta onto the frozen library of mechanisms to obtain cosine alignment scores (Fig. 5f). The resulting rankings prioritize partners whose directions best align with the desired move; top candidates in our analysis include α-PD-1/IL-15^38–41^, α-PD-1/IFN-gamma^42–44^, α-PD-1/IL-2^45,46^, and α-PD-1/IFN-alpha^47^ (Fig. 5g), all of which, except α-PD-1/IL-34 have literature support as α-PD-1 augmenters. Head-to-head on the identical harmonized data foundation, CellFlow^23^ produced a different ordering with generally weaker concordance to literature reported synergies^48–50^ (Fig. 5h).

To interrogate how TRAILBLAZER’s predictions align with the biology of the tumor immune microenvironment, we coupled the simulator to a multicellular phenotype classifier that produces cell-importance readouts and then organized both mechanisms and patients in the space defined by these readouts. The classifier aggregates information across hundreds of cells with an attention-style pooling layer; the learned weights provide a per-sample map of which cell populations are the most important to the phenotype of interest (referred here as importance), predicted probability of the classifier is used to assess likeness to the desired phenotype (referred here as responsiveness) (Fig. 6a). We have observed that embeddings produced with set transformers, as compared to single cell model embeddings produced by Geneformer^51^, scGPT^9^ and scVI^52^ yield higher F1 score of multilabel phenotypic state classification assessed on CZ CELLxGENE Discover human disease datasets^53,54^(Fig. 6b). Aggregating importance weights by annotated cell type yields cell importance scores for each patient and condition (Fig. 6c and Ext. Fig. 5). Importance scores are stable across bootstrap resampling and seeds, and their principal axes correspond to canonical immunobiology, providing an interpretable link from single-cell predictions to system-level readouts.

**Fig. 6.**
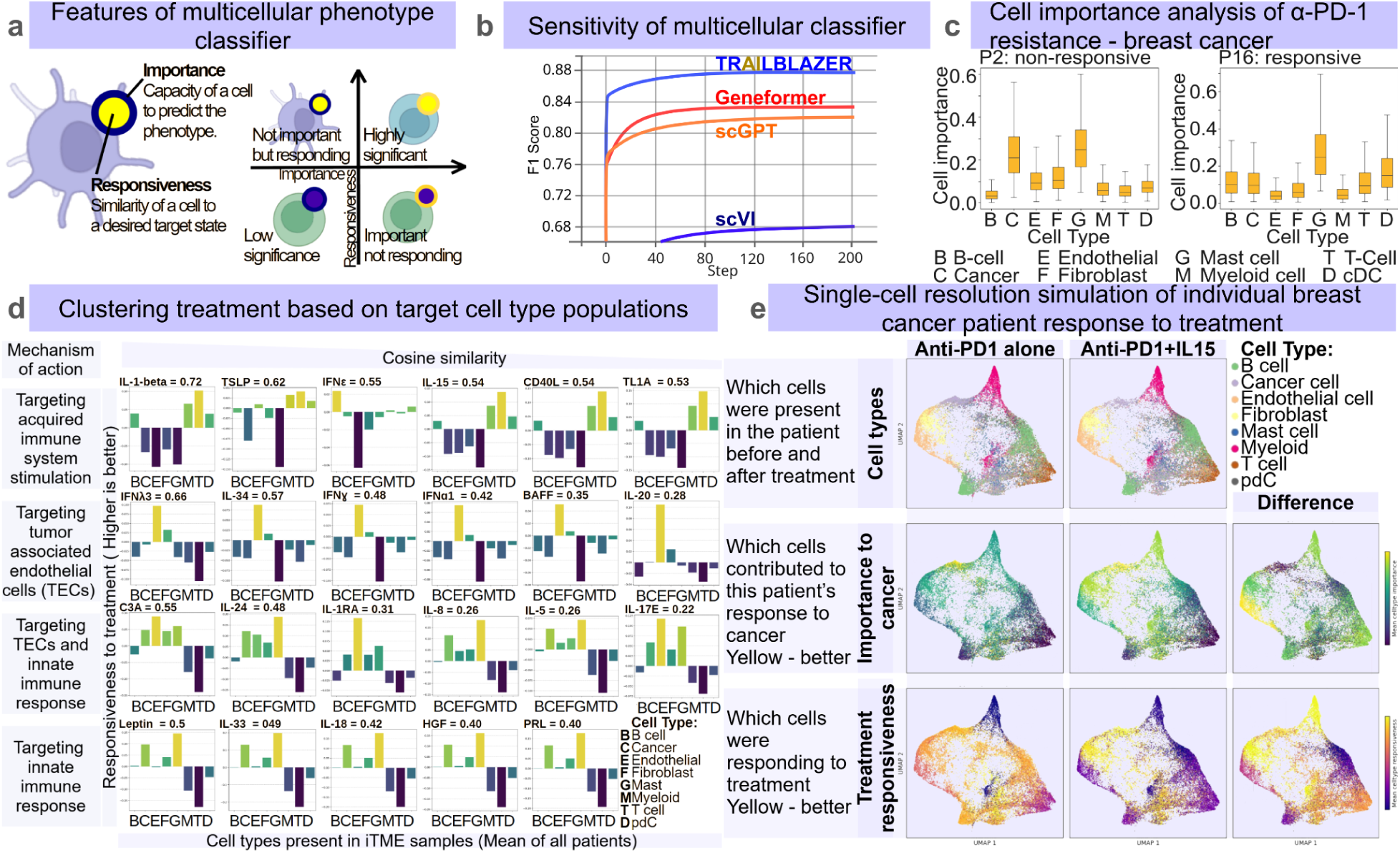
TRAILBLAZER’s cell attention can be used to aid granular analysis of diseases and treatments. **a**, The interpretability of the model predictions is afforded by multicellular phenotype classifier, trained to recognize desirable and undesirable cell states (e.g. responsiveness to treatment). It is capable of assessing cells’ state contribution to a disease/phenotype (cell importance), and it measures cells’ similarity to a desired state after treatment (responsiveness). **b**, The TRAILBLAZER phenotype classifier’s multicellular architecture outperforms the single-cell architectures of state-of-the-art models (Geneformer, scGPT, scVI). Models were trained on multilabel disease classification using patients’ scRNA-seq data, then evaluated on unseen patients. Performance was measured as F1 score of disease classification across training epochs. **c**, An example of TRAILBLAZER’s inferred importance of each cell type for the naive breast cancer patients, for a patient resistant to anti-PD-1 (left), and responsive (right). **d**, TRAILBLAZER AI predicted treatments improving anti-PD-1 patient responsiveness clustered by target cell populations. The average treatment responsiveness of each cell type is calculated across all patients (N=9). Treatments are clustered into four groups based on their predicted mechanism of action and are sorted from left to right based on the overall cosine similarity of cell states to positive patient outcomes. **e**, TRAILBLAZER AI models the contribution of each cell to anti-PD-1 patient responsiveness, an example of anti-PD-1/IL-15 effect on tumor micro-environment. A detailed view with single-cell resolution shows cells present before (1st column, top row) and after treatment (2nd column, top row), the importance of each cell to the state (middle row), and the treatment responsiveness of each cell (bottom row).

We group perturbations into mechanism families by how they redistribute cell-importance across populations. For each agent we simulate the counterfactual post-treatment state, recompute cell-importance maps, and take the delta relative to the pre-treatment baseline; clustering these deltas across agents reveals distinct mechanism clusters. Because clusters are defined by population-level deltas, they also provide an actionable summary of how each candidate therapy is expected to reshape the microenvironment. Analysis identified four clusters of treatments augmenting anti-PD-1, acting through: acquired immune response cells; innate immune response cells; tumor associated endothelial cells (TECs); TECs with innate immune response (Fig. 6d). These clusters coincide with neighborhoods in the intervention embedding library, and the top-ranked partners in our combination screens tend to fall into the acquired-immunity family, consistent with literature^38–47^.

In Fig. 6e we illustrate patient response to α-PD-1 and α-PD-1/IL-15 combination at a single-cell resolution by projecting pre- and post-simulation cells onto a common cell set UMAP and overlaying cell-importance and delta of responsiveness, highlighting which compartments shift and how those shifts. Together, these panels show which populations drive the predicted effect, which treatments act through similar pathways, and which patients are likely to share a therapeutic strategy.

## DISCUSSION

Our results argue that multicellular context is the right abstraction for modeling immune perturbations at single-cell resolution. Empirically, when we increase the number of cells per set at training time, while holding the total number of cells per step constant, distributional reconstruction improves monotonically (Fig. 2c,d; Ext. Fig. 2c). Likewise, the perturbation segmentation encoder collapses when sets are too small because intra- and inter-state distances become indistinguishable (Fig. 2e–h), indicating that a minimum cell population context is required to form stable state centroids. These findings are consistent with a view of tissues as dynamical systems in which collective constraints stabilize phenotypes even when any one cell is noisy; we stipulate that a multicellular model can extract and reuse those constraints, improving generalization.

Multicellular encoding changes what the model can represent. Single-cell approaches lack an explicit interface for simulating coordinated shifts in population structure. Used here set encoder, coupled with balanced sampling and donor matching, forces the model to learn within cell-type shifts and population-level couplings rather than shortcutting through changes in cell composition. This design isolates the causal signal of interest, how a given mechanism moves a given donor’s populations in gene-expression space. In practice, the encoder’s near-linear scaling (Ext. Fig. 2a,b) makes such modeling feasible at clinically relevant cell set sizes.

Equally important is the organization of the latent space. We shape the geometry so that healthy states cluster near a common reference and interventions correspond to calibrated directions whose angles encode mechanism and whose radii reflect magnitude. This geometry functions as an interface: it lets us add or subtract interventions (x→x+g, y→y−g) to simulate counterfactuals; project a desired shift onto a library of mechanisms to obtain patient-specific rankings; and compose directions to model combinations. The qualitative effect of shaping is a reorientation from donor-dominated manifolds to mechanism-dominated clusters with donor substructure (Fig. 3b–e); quantitatively, rediscovery improves from approximately random (top-45) to top-5 (Fig. 3f), and enables zero-shot generalization to an unseen treatment with further improvements from few-shot supervision (Fig. 4). There is, however, a modest trade-off in reconstruction tightness (Fig. 3g), but the gain in semantic structure, projection and composition, enables use cases that pure reconstruction struggles to support.

We postulate that a unifying explanation for these behaviors is a generalization hypothesis, according to which immune tissues operate with a relatively small repertoire of multicellular behaviors (“archetypes”), and these behaviors can be captured as directions in a shaped latent space. Once geometry is imposed, mechanism clusters emerge (Fig. 3b–e), consistent with shared response programs that recur across individuals. The mechanism library learned independently of reconstruction is low-dimensional and concentrated, per-intervention embeddings fit by von Mises–Fisher distributions have tight concentration parameters, and a small set of directions explains much of the variance in donor×perturbation latents. Under the generalization hypothesis, shaping improves rediscovery because directions align with “archetype” mechanisms, which also explains why zero-shot performance approaches few-shot as new conditions are composed rather than memorized. It also suggests natural extensions: dosing as scaling of directions, scheduling as sequences, multimodality as conditioning of the decoder, and cross-domain transfer as alignment of mechanism bases across tissues or species.

There are, however, limitations and open directions. First, shaping introduces a modest trade-off between reconstruction tightness and semantic structure, per-cell-type energy distance increases slightly under shaping (Fig. 3g), even as rediscovery and zero-/few-shot performance improve. Relatedly, while negative-binomial and zero-inflated NB heads preserve mean–variance relationships and zero inflation, they can over-smooth low-count regimes; presumably conditional diffusion decoders could deliver higher per-cell fidelity at the cost of inference speed and are a natural next step. Second, results depend on upstream harmonization and the chosen gene space. Although our harmonizer removes depth and batch confounders while preserving responder signal, strong shifts in chemistry or tissue composition may require retraining adapters. Furthermore, restricting gene sets to highly variable genes simplifies modeling at the risk of discarding predictive rare markers.

Third, combination modeling assumes approximate additivity of directions; while this captures many cases, true biological interactions can be sub- or super-additive, dose- and time-dependent, or order-sensitive. To address these issues we propose extending the interface to include coefficients (dose), sequences (schedules), and adding interaction terms. Fourth, balanced sampling and donor matching in the dataloader isolate within-cell-type shifts and prevent composition shortcuts during training; this is a feature for causal inference but a limitation when compositional changes are themselves part of the effect size (e.g., cell proliferation, or recruitment). A practical compromise is to expose composition-aware readouts downstream while keeping the encoder trained on balanced sets, or to add a dedicated head that models compositional dynamics alongside within-cell-type shifts.

Fifth, the present work is unimodal and assumes well-annotated cell types. In microenvironments shaped by protein signaling, spatial proximity, or chromatin state, adding CITE-seq (protein), spatial RNA, or scATAC conditioning to the decoder should improve context awareness and interpretability. The same set-based encoder can accommodate neighborhood priors for spatial context or cross-modal adapters for RNA-protein alignment. Sixth, cross-species translation remains a key goal for preclinical screening. Our mechanism-library view naturally extends to a bi-encoder with species adapters and shared geometry; aligning mechanism bases across species would enable predicting human-like effects from mouse data without exhaustive human screening.

Finally, the proposed architecture is specifically designed for multicellular, heterogeneous systems that more closely resemble physiological tissue environments. In contrast, the majority of currently available perturbation datasets are derived from isolated cell lines, which lack the intercellular communication and contextual signaling present in vivo. Such reductionist systems are therefore suboptimal for training models intended to capture multicellular perturbation dynamics. Conversely, in the absence of cellular crosstalk and microenvironmental complexity, multicellular modeling frameworks offer little advantage over single-cell modeling for predicting cell line behavior.

Despite these limitations, we are convinced that the solutions presented are a significant step toward enabling virtual trials that are both plausible and actionable, helping match the right treatment to the right patients.

## EXTENDED FIGURES

**Ext. Fig. 1.**
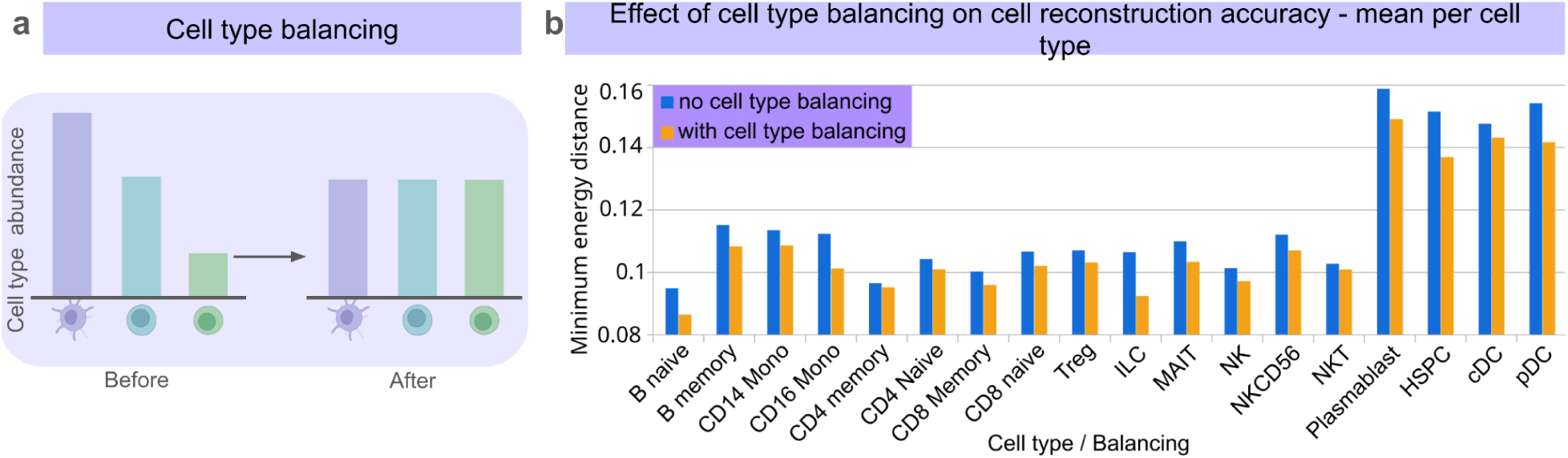
Cell type balancing improves training accuracy. **a**, Graphical abstract of cell type balancing. **b**, Minimum energy distance for network with and without cell type balancing calculated for each gene, averaged across 25 cells per cell type, for reconstruction of scRNA-seq of unseen donors (N=3), trained on sets of 500 cells. All measurements use the PARSE PBMCs scRNA-seq dataset.

**Ext. Fig. 2.**
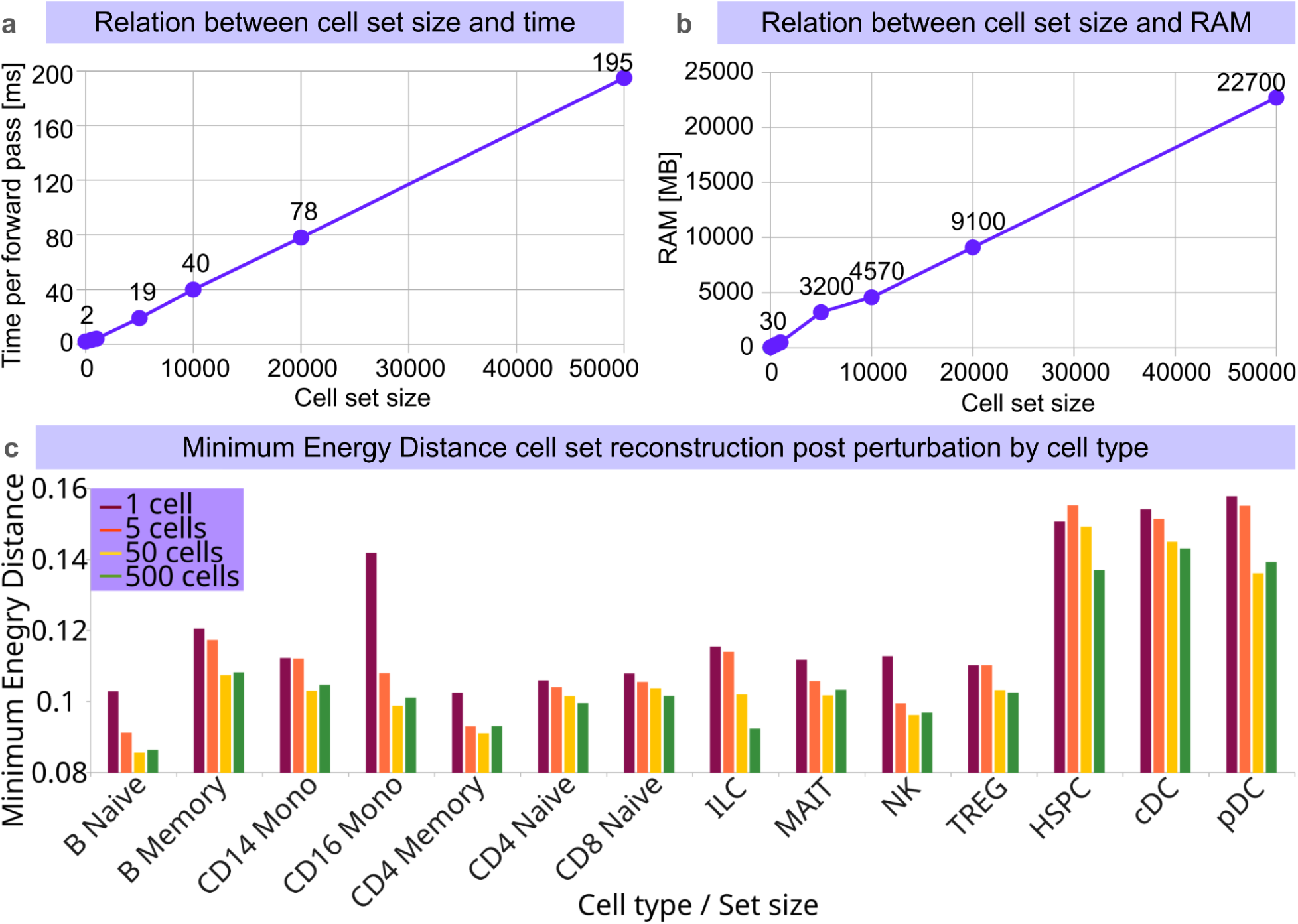
Set transformers as attention-based permutation-invariant encoders for cell sets. **a**, Relation between cell set size and time necessary to complete a single forward pass. **b**, Relation between cell set size and RAM use. **c**, Energy distance calculated for each gene per cell type, averaged across 25 cells per cell type, for reconstruction of scRNA-seq of unseen donors (N=3), calculated every 100 step of the training on sets of 1, 5, 50 and 500 cells, for 90 treatments and 9 donors. In all runs, the network sees exactly 1000 cells in each forward pass, independent of cell set size. Latent shaping was not applied. All measurements use PARSE PBMCs scRNA-seq dataset, where the genes were subset to a group of 5634 HVG genes.

**Extended Fig. 3.**
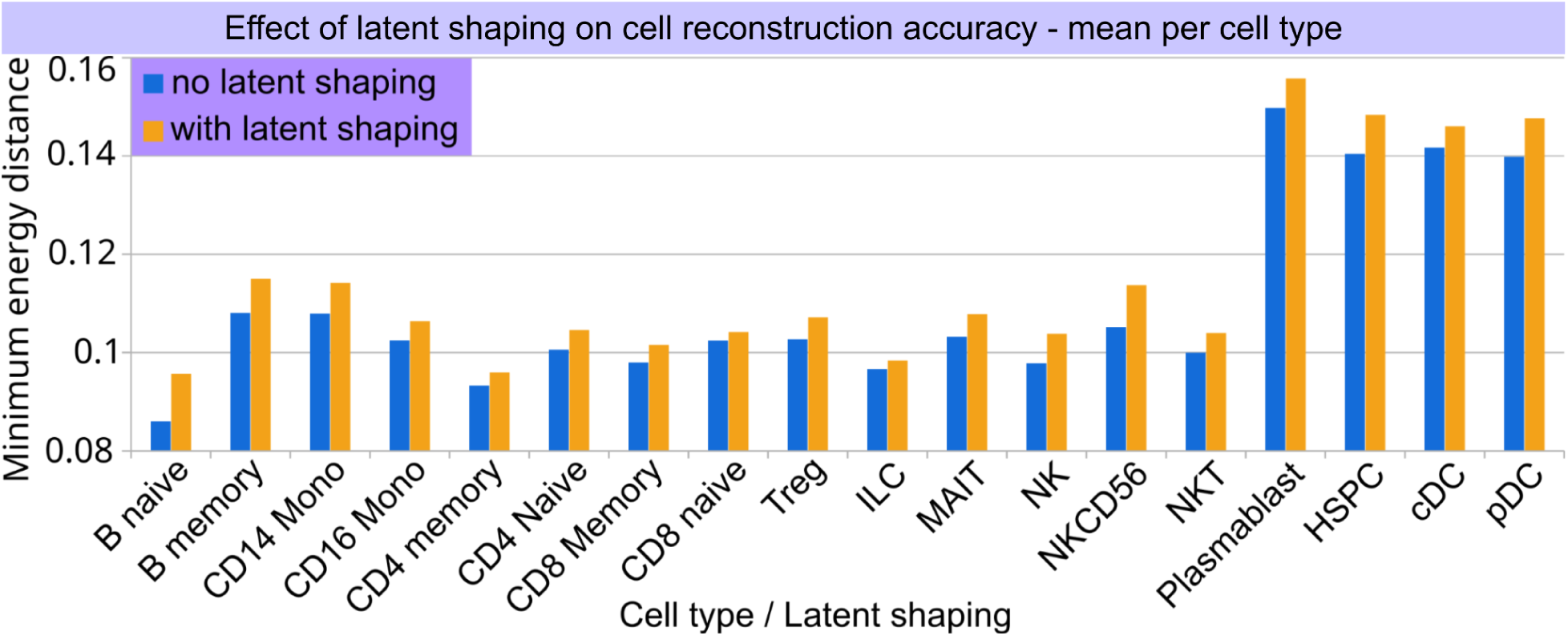
Application of latent space shaping using biological priors. Minimum energy distance per cell type for network without and with latent shaping calculated for each gene, averaged across 25 cells per cell type, for reconstruction of scRNA-seq of unseen donors (N=3), trained on sets of 500 cells. All measurements use PARSE PBMCs scRNA-seq dataset.

**Extended Fig. 4.**
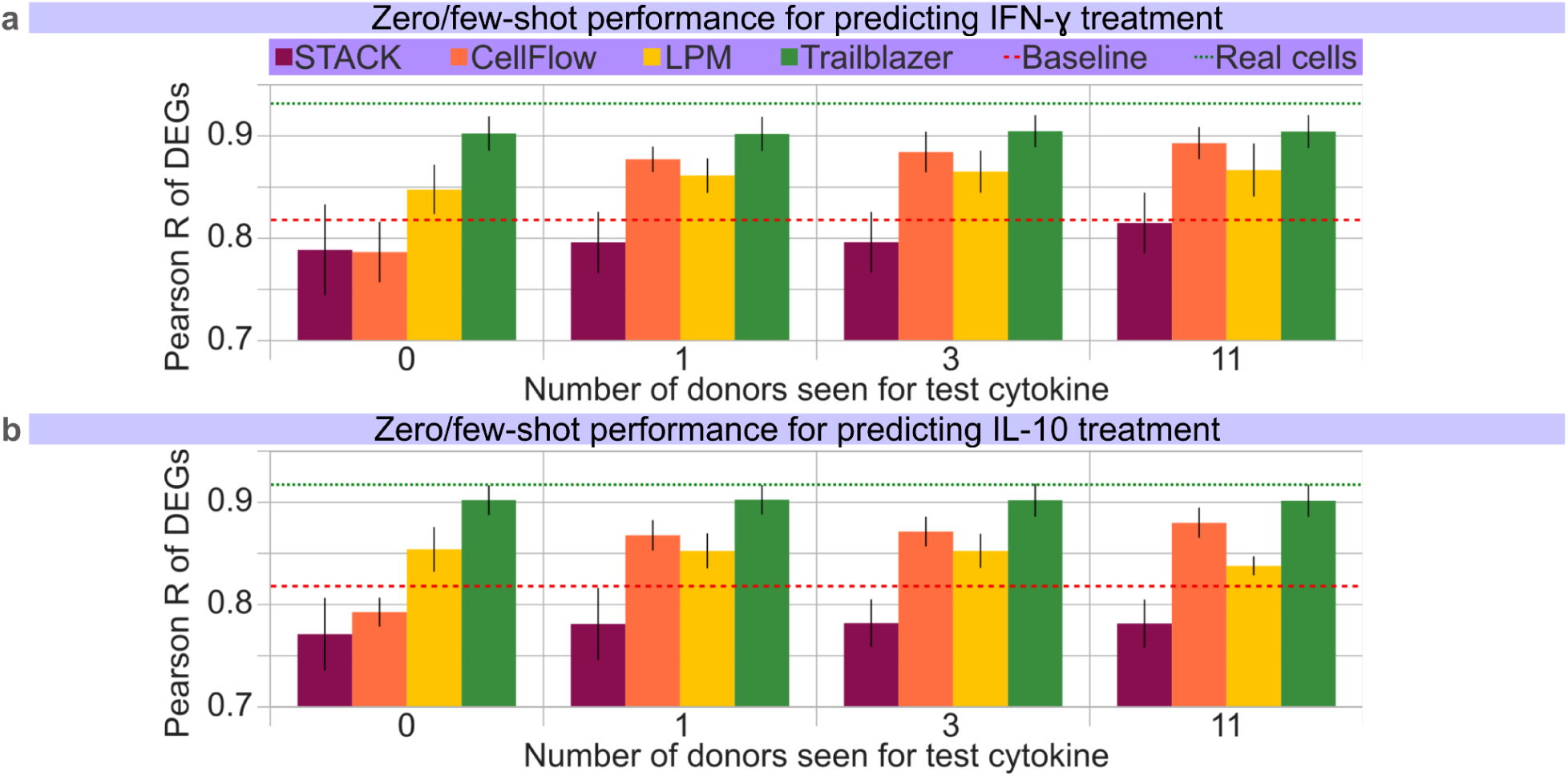
TRAILBLAZER’s architecture exhibits superior performance for both zero-shot and few-shot treatment outcome prediction. **a**, Post-treatment reconstruction Pearson R score for top 100 differentially expressed genes in a task of predicting post-treatment scRNA-seq state of the withheld donor for networks trained on 0, 1, 3 and 11 donor examples of the test treatment. Baseline control indicates no change from pretreatment state, real cells denote ground truth, sampled in the same fashion as predictions. Networks are trained on a common dataset of ex vivo 90 treatments of Peripheral Blood Mononuclear Cells from 12 donors. Measurements are means of 100 samples and their standard deviation. Comparison of STACK, LPM, CellFlow, and Trailblazer models performance for IFN-ɣ; treatment prediction; **b**, IL-10. All measurements use the PARSE PBMCs scRNA-seq dataset.

**Extended Fig. 5.**
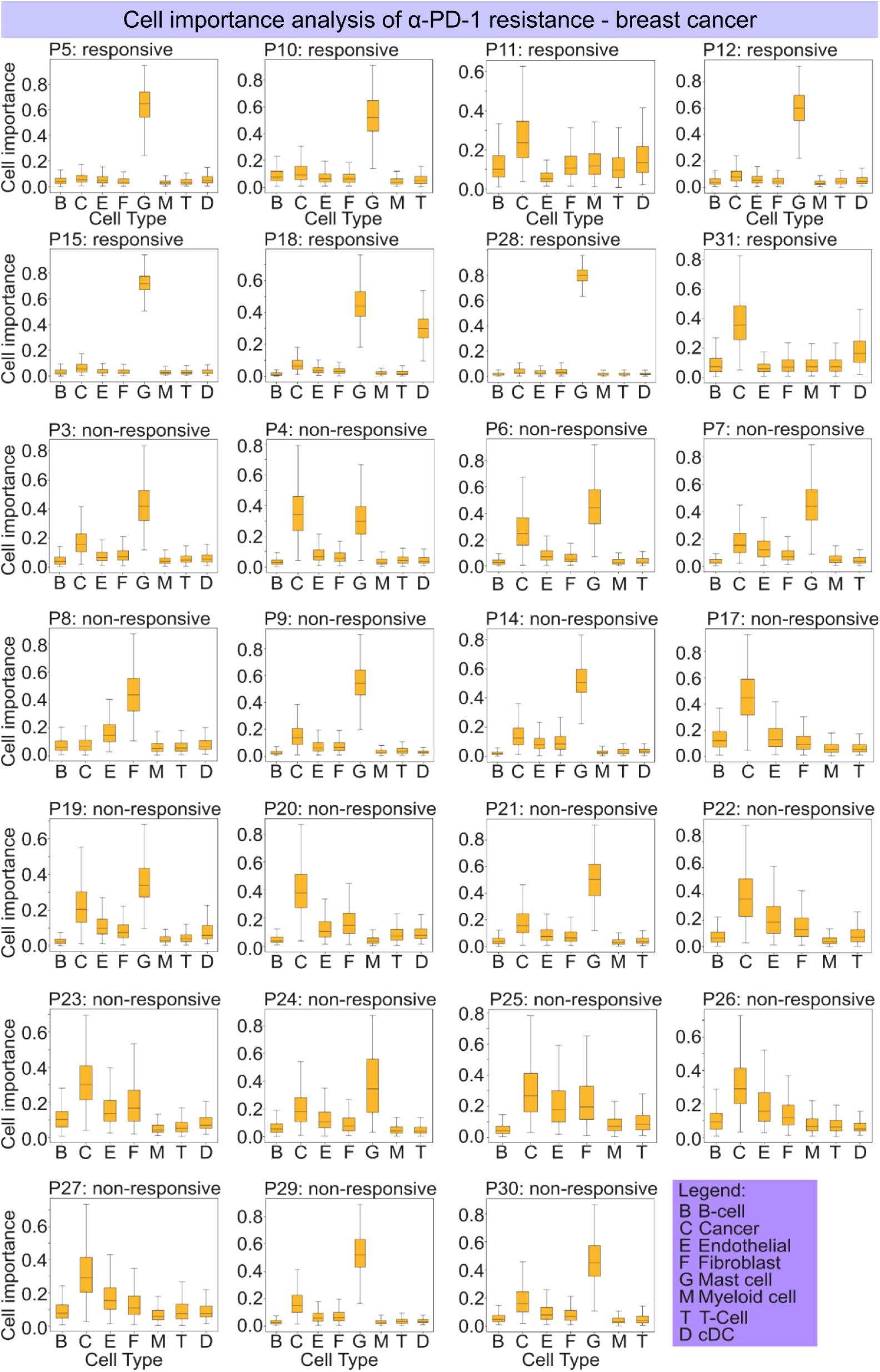
TRAILBLAZER’s cell attention can be used to aid granular analysis of diseases and treatments. TRAILBLAZER’s inferred importance of each cell type for the naive breast cancer patients, for a patient that was treated with anti-PD-1. Based on dataset, cohort A of “A single-cell map of intratumoral changes during anti-PD1 treatment of patients with breast cancer”.

## METHODS

### Dataset construction

The dataset used to train and validate the models was based off of the data corpus containing PBMCs perturbations created by PARSE^30^. To facilitate rapid architecture development, the dataset was first subset to 1M cells in such a way that there is maximum variability across cell types: if there are less cells present than needed for a given cell type to achieve a uniform distribution, then all of them are taken. This way, rare cell types are overrepresented while abundant cell types are underrepresented in the subset. Subsetting had minimal impact on final results as compared to full set.

Only cells with less than 10% mitochondrial content were kept. To filter genes, highly variable genes (HVGs) were identified using Scanpy^55^ ver 1.12. After lognormalizing the counts, genes were required to have an average expression between 0.0125 and 3 to exclude extremely lowly and highly expressed genes. Among those genes, we further retained only genes with dispersion greater than 0.25. From here we obtained a total of 5301 genes. We further added an extra set of about ∼300 gene markers that we considered important cell type identifiers, getting a final set of 5634 genes (Supplementary data 1).

Finally, we performed a filtering step by keeping cells that have at least 100 counts, ending up with a total of 998657 cells.

For testing and downstream analysis, a dataset comprised of naive and post-treatment breast cancer patients treated with anti-PD1^33^, along with CELLxGENE^53^ were also part of the data used for training and validation during the elaboration of our experiments, processed in similar fashion to PARSE PBMCs dataset.

### Dataloader design

To facilitate model training, we use a purpose-built dataloader that stratifies by donor, state, and cell type; enforces per–cell-type quotas within each group; and augments scarce populations to avoid mode collapse on abundant cell types. The loader is disk-backed and chunked to draw fairly from tens of millions of cells without exhausting RAM, with an in-memory cache to maintain throughput. Every batch returns permutation-invariant views of the paired sets (control and perturbed), with cell order randomized on each draw and masks/padding handling variable set sizes. To attribute gains from larger sets to genuine multicellular reasoning rather than averaging artifacts, we fix the product of batch size and set size (e.g., 1,000 cells per forward pass) across conditions; this constraint is enforced directly in the loader. When latent arithmetic is required, the same batch is accompanied by the corresponding intervention identifier so that forward (control → apply intervention) and reverse (perturbed → remove intervention) passes can be formed without additional I/O. To aid the pipeline reproducibility we use deterministic seeding and on-the-fly label projections (for post-hoc cell-type stratification).

The network expects two groups of cells for each training iteration: a control group, and its corresponding perturbed state.

Let *D* denote the full set of cells available for training. Each cell c ∈ *D* is associated with a set of categorical covariates, including dataset *d*(c), donor *s*(c), perturbation condition *p*(c), and cell type *t*(c).

The objective of the dataloader is to generate pairs of cell groups (*G*_ctrl_, *G*_pert_), each of fixed size *N*_cg_, while minimizing sampling bias across these covariates. In particular, the sampling procedure is designed to avoid over-representing dominant datasets, donors, perturbations, or cell types.

As a preprocessing step, cells are grouped and cached according to their full covariate tuple (*d, s, p, t*). Formally, for each combination of covariates we define the index set

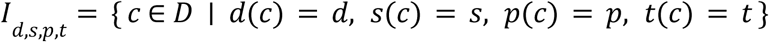

To allow the dataloader to sample covariate combinations efficiently during training without repeated filtering over the full dataset, we implemented hierarchical mappings that associate each dataset with its available perturbations, each dataset-perturbation pair with its donors, and each dataset-donor-perturbation tuple with its observed cell types.

To mitigate biases arising from uneven cell counts across experimental conditions, for each training iteration, a covariate tuple is sampled uniformly in a sequential manner. First, a dataset *d* is selected uniformly from the set of datasets. Conditioned on *d*, a donor *s* is selected uniformly from the donors available in that dataset. Finally, a perturbation *p* is selected uniformly from the perturbations observed for the chosen dataset-donor pair.

Given a sampled tuple (*d, s, p*), the corresponding control condition is defined as the non-perturbed state with the same covariates, i.e. (*d, s, control*).

Let

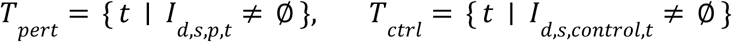

Then, only cell types present in both conditions are considered for group construction. The matched cell type set is therefore given by

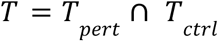

This restriction ensures that control and perturbed groups are comparable at the cell type level. To construct cell groups of fixed size *N*_cg_, a set of non-negative integer counts {n_t_}_t∈T_ is sampled uniformly subject to the constraint

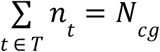

This corresponds to sampling uniformly from the discrete simplex defined by the matched cell types. As a result, no particular cell type is systematically favored across training iterations, and the relative abundance of cell types varies randomly while remaining identical between the control and perturbed groups.

To guarantee that both groups share the same per-cell-type composition, for each cell type t ∈ *T*, exactly n_t_ cells are sampled independently for the perturbed and control conditions from the corresponding index sets *I*_d,s,p,t_ and *I*_d,s,control,t_, respectively. This guarantees that both groups share the same per-cell-type composition.

In cases where the number of available cells for a given covariate tuple and cell type is smaller than the required count n_t_, data augmentation is applied. Specifically, all available cells of that type are first included, and the remaining cells are obtained by sampling with replacement from the available set. For each duplicated cell, a new expression profile is generated by sampling each gene expression value independently from a Poisson distribution with rate parameter equal to the original expression value,

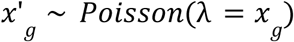

where x_g_ denotes the original count for gene g.

The final output of the dataloader for a given iteration is then our desired cell group pair (*G*_ctrl_, *G*_pert_), each of size *N*_cg_, with identical cell type composition and matched dataset and donor covariates.

### The TRAILBLAZER model

The TRAILBLAZER model is designed to operate on groups of cells rather than individual cells, and to explicitly disentangle intrinsic cell identity from perturbation-driven effects. Given paired control and perturbed cell groups sampled as described above, the model learns a latent representation of cell identity and a latent representation of perturbation effects. For decoding back to gene counts we use a negative-binomial or zero-inflated negative-binomial parameterization to reconstruct full expression vectors. Decoder layers are modulated with feature-wise linear modulation (FiLM)^56^, allowing us to condition on context (e.g., perturbation embedding, dataset/cell-line) so that systematic “style” is re-injected at decode time without contaminating the shared latent geometry.

Let *X* ∈ ℝ^Ncg×G^ and *Y* ∈ ℝ^Ncg×G^ denote control and perturbed cell groups, respectively, where *N*_cg_ is the number of cells per group, and *G* is the number of genes. In addition, each batch is associated with a perturbation embedding g ∈ ℝ^1^ ^×^ ^dz^, derived from an external encoder described in the next section.

Cell identity is modeled using a shared encoder E_cell_ applied independently to control and perturbed groups. Given a cell group *X*, the encoder produces a latent representation

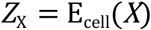

where *Z*_X_ ∈ ℝ^Ncg^ ^×^ ^dz^ and *d*_z_ denotes the latent dimensionality.

The encoder first projects gene expression vectors into a hidden feature space using a pointwise linear transformation to capture per-cell features. Then, to capture global group-level context, the projected representations are processed using a Set Transformer architecture. These two steps produce two complementary components: cell-specific embeddings capturing local variation, and a global group embedding summarizing shared structure across the group. The global embedding is broadcast back to all cells and concatenated with the cell-specific features, yielding a representation that jointly encodes individual cell identity and group context.

The contextualized features are then passed through an output projection to parametrize a Gaussian latent space. The model then outputs mean μ and a log-variance log σ for each of the parameters, as is standard in variational autoencoders. Finally, latent variables are sampled using the reparametrization trick,

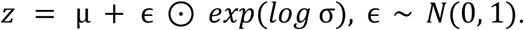

The same encoder is shared for both control and perturbed groups, yielding latent representations *Z*_X_ and *Z*_Y_, respectively.

Perturbation effects are modeled as additive shifts in latent space. After broadcasting the perturbation embedding *g* to match the cell dimension, the model constructs four latent configurations: *Z*_X_ representing the control identity, *Z*_Y_ representing the perturbation identity, (*Z*_X_+g) representing the perturbed state derived from control cells, and (*Z*_Y_ **-** g) representing the control state derived from perturbed cells. This arithmetic structure encourages the latent space to organize such that perturbation effects are approximately linear and compositional.

Gene expression reconstruction is performed using a shared zero-inflated negative binomial decoder D_ZINB_, which maps latent representations back to gene space. Given a latent vector *Z*, the decoder outputs parameters of a zero-inflated negative binomial distribution:

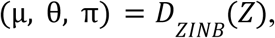

where,

μ ∈ ℝ ^G^ denotes the mean expression, θ ∈ ℝ ^G^ denotes the inverse dispersion, and π ∈ ℝ^G^ the zero-inflation logits.

The decoder consists of a multi-layer perceptron with ELU activations and dropout regularization. Positivity of μ and θ is enforced via a softplus transformation. θ is clamped to avoid extreme values and favor numerical stability. This decoder is applied in each training iteration to all four latent configurations previously described.

### Training losses

The model is trained end-to-end using a composite objective that enforces accurate reconstruction of observed cell groups, consistency under perturbation arithmetic in latent space, and distributional alignment between predicted and observed gene expression profiles. Since both inputs and outputs are treated as unordered sets of cells, all losses are defined in a permutation-aware manner.

Decoder outputs correspond to unordered sets of reconstructed cells. To compare predicted and observed cell groups at the cell level, we first compute an optimal matching between predicted and target cells using a Chamfer-style assignment based on reconstructed mean expression profiles. This produces a permutation of predicted cells that minimizes pairwise distances to the target set. All decoder output parameters μ, θ and π are reordered according to this assignment before loss computation.

Gene expression counts are modeled using a zero-inflated negative binomial (ZINB) distribution. For each reconstructed cell *i* and gene *j*, the decoder predicts parameters (µ_ij_, θ_ij_, π_ij_), defining a ZINB likelihood

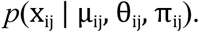

We apply the ZINB negative log-likelihood to four reconstruction pathways: identity reconstructions (*X* → *X*) and (*Y* → *Y*), and cross-condition reconstructions (*X* → *Y*) and (*Y* → *X*). We label these reconstruction losses as

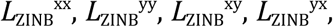

each weighted by a corresponding scalar coefficient. Identity reconstructions encourage faithful encoding and decoding of observed states, while cross-condition reconstructions enforce that latent perturbation arithmetic remains consistent.

While ZINB reconstruction losses operate at the individual cell level, we additionally enforce group-level consistency between predicted distributions. For each reconstructed group, we compute the mean ZINB parameters across cells. We then penalize discrepancies between group-level parameters of inverse reconstructions and their corresponding identity reconstructions. Specifically, we enforce that the mean parameters of (*Y* → *X*) match those of (*X* → *X*), and that the mean parameters of (*X* → *Y*) match those of (*Y* → *Y*).

Losses on μ and θ are computed in log space to stabilize optimization across genes with widely varying expression levels, while losses on π are computed directly, obtaining

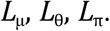

Gradients are stopped on identity reconstructions, making them fixed references.

We also impose explicit geometric constraints on the latent space to encourage perturbation effects to be well aligned, additive, and metrically consistent across cell groups.

Let *Z*_X_ and *Z*_Y_ denote the latent representations of control and perturbed cell groups, respectively, and let *g* denote the corresponding perturbation embedding. To obtain group level summaries, we compute mean latent vectors

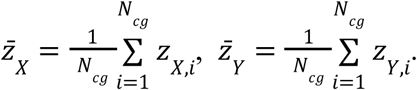

To encourage perturbations to correspond to consistent directions in latent space, we enforce alignment between the displacement vector z̄_Y_ - z̄_X_ and the perturbation embedding *g*. This is implemented using a cosine similarity loss,

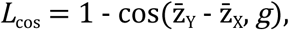

which promotes a shared geometric interpretation of perturbations across different cell groups.

To further structure the latent space, we apply norm based constraints to the group level latent representations. First, control group representations are encouraged to lie near the origin,

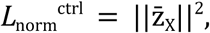

and second, the norm of perturbed group representations is encouraged to match the unit norm of the perturbation embedding,

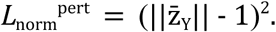

Together, these constraints encourage perturbed states to lie around the surface of a hypersphere of unit radius, while controls lie close to the origin.

The latent metric losses are combined with the reconstruction and distributional objectives described above, each weighted by a corresponding coefficient, obtaining the total loss as a weighted sum of each of the individual losses:

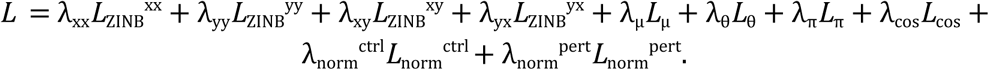

### Training strategy and metrics

Cell sets of fixed size *N*_cg_ = 500 were used throughout training. We choose this set size based on the dataset’s limitations (small sample set sizes) and to avoid the need for excessive cell augmentation when uniformly sampling across all cell types and perturbations.

Directly optimizing the full objective by combining reconstruction losses and latent metric regularization from the start was found to lead to suboptimal convergence, with the model frequently becoming trapped in poor local minima. To mitigate this behavior, we adopt a staged training strategy in which different components of the loss are progressively introduced.

In the first stage, we train reconstruction and apply only the radial term for controls, allowing the encoder–decoder to settle while anchoring the healthy reference. During that stage, we place greater emphasis on the reconstruction loss *L*_ZINB_ ^yy^ by setting λ ≫ λ.

In the second stage, we introduce the angular (cosine) alignment so that perturbation directions behave consistently across donors and tissues; cross-condition reconstruction losses *L*_ZINB_ ^xy^ and *L*_ZINB_ ^yx^ are enabled, allowing the model to learn mappings between control and perturbed latent representations. Training is continued until reconstruction losses and validation metrics reach a stable plateau.

In the final stage, latent metric regularization is introduced to shape the geometry of the latent space. We first enable the directional alignment loss *L*_cos_ together with the control norm regularization term *L*_norm_^ctrl^ to pull control states towards the center of the hypersphere. Once these terms have stabilized, the perturbation norm regularization *L*_norm_^pert^ is activated. For numerical stability and to prevent over-constraining the latent space, all norm-based losses are assigned relatively small weights throughout training.

### The mechanism segmentation model & dataloading strategy

The data loading strategy closely follows that described in the previous section, with the key difference that, instead of sampling pairs of cell groups, we first sample *N* treatments uniformly at random. Then, for each treatment, *M* cell groups are drawn following the same covariate aware strategy as before. This yields a final tensor of dimension N × M × *N*_cg_ × G.

This model, much like TRAILBLAZER, is also designed to work on unordered sets. To obtain a fixed dimensional embedding from an unordered collection of input observations, we employ a Set Transformer based encoder. Given an input set

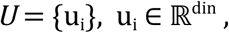

the goal of the encoder is to produce a single embedding vector

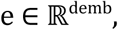

that is invariant to permutations of the elements in *U*.

Each element u_i_is first projected independently into a shared hidden space using a linear transformation, producing a set of hidden representations. This resulting set is then processed by a Set Transformer module configured to produce a single output token. This pooling operation aggregates information across all set elements while remaining invariant to permutations of the input set.

The pooled representation is subsequently passed through a linear projection and a leaky ReLU activation to obtain embedding. Finally, the embedding is normalized to unit length, which constrains all embeddings to lie on the unit hypersphere and facilitates the use of cosine similarity as a comparison metric.

This final representation is denoted as *g* throughout the paper. The embedding dimensionality was fixed to 16, as empirical experiments indicated that this size offers an adequate trade-off between representational capacity and model complexity for the 90 treatments in the PARSE dataset.

### Inference

At inference we take a patient’s harmonized control sample and a perturbation to simulate the counterfactual post-treatment state. We sample one or more control sets from the patient, encode them to per-cell latents x, add the unit-norm mechanism vector g for the chosen perturbation to obtain x+g, optionally scaled to reflect dose or exposure, and decoded to full gene-count vectors using the conditioned likelihood head, where FiLM layers ensure the decoder matches the patient’s dataset/cell-line style. To increase robustness we sample multiple latent realizations per intervention from a compact distribution fitted during training (e.g., a von Mises–Fisher over mechanism latents) and average summaries across draws. The resulting counterfactual single-cell sets are analyzed per cell type to compute expression deltas (perturbed minus control), visualize distributions (UMAPs, violin plots), and quantify distributional shifts with energy distance.

To prioritize treatments, we estimate the delta needed to move the patient toward a desired target state (e.g., a healthier or responder-like state) and project that delta onto the frozen library of mechanism vectors. The cosine between the desired delta and each library vector yields a patient-specific ranking, with higher alignment indicating a better match; this rediscovery/ranking signal is the same geometric quantity used during training and correlates with zero-/few-shot performance. For combinations we compose directions additively, g₁+g₂(+⋯), optionally scaling coefficients to reflect dose or timing; small grids or greedy searches over coefficients suffice in practice, and the normed geometry stabilizes composition across donors. Rankings are reported globally and per cell type or cell set, together with contribution heat maps that indicate which populations drive the predicted shift.

To translate molecular predictions into clinical readouts, we coupled counterfactual state predictions to phenotype classifiers. For anti-PD-1 response prediction, a multicellular responder/non-responder classifier was applied to the simulated post-treatment cells to estimate the probability that the patient will respond to anti-PD-1. All inference routines honor the permutation-invariant setup used in training (randomized group order, fixed batch size×set size), and we report uncertainty via bootstrap confidence intervals over set/draws.

### Virtual patient treatment

To simulate in silico anti-PD-1 treatment and assess predicted responsiveness, the anti-PD-1 breast cancer dataset^33,34^, cohort A, was combined with the previously described 1M PARSE subset^30^. The anti-PD-1 dataset was first preprocessed using the same pipeline applied to PARSE, with the additional exclusion of cells lacking expansion metadata. To align with the 5634 genes present in PARSE, missing genes were imputed with zeros.

Given the substantial batch effects present between datasets attributable to differences in sequencing technology, experimental conditions, and sample collection procedures, the anti-PD1 dataset was harmonized using our proprietary harmonizer architecture prior to integration with PARSE. This step was taken to align the technical variation profile of the anti-PD1 dataset with that of PARSE, ensuring that residual differences between datasets reflect genuine biological signals rather than technical artifacts.

The mechanism segmentation network was then trained incorporating, alongside the perturbed states from PARSE, the post-perturbation profiles of responding patients from the anti-PD-1 dataset. These profiles define the anti-PD-1 embedding within the segmentation network.

The main TRAILBLAZER network was subsequently trained using PARSE, and both responding and non-responding patients from the anti-PD-1 dataset, allowing the network to learn that applying the anti-PD-1 embedding to naive patient states can yield either a responder or non-responder outcome depending on the individual. During the latent shaping stage of TRAILBLAZER training, naive states from the anti-PD-1 dataset were excluded from being pulled towards the center of the latent space, as these states are not considered representative of a healthy baseline.

Post-perturbation states were inferred by applying the trained TRAILBLAZER network to held-out naive patients from the anti-PD-1 dataset, generating approximately 100 synthetic samples of 500 cells per patient. The resulting synthetic cells were then evaluated using the multicellular anti-PD-1 responsiveness classifier, defined as TCR clonotype expansion, whereas TCR-seq data was not used for training or inference, and predicted responsiveness scores were summarized as violin plots.

### Virtual drug treatments

The TRAILBLAZER network trained on the combined PARSE and anti-PD-1 dataset was additionally used to generate a ranking of drug candidates predicted to augment anti-PD-1 response. Pre and post-treatment samples from responding patients in the anti-PD-1 dataset (approximately 100 samples per donor) were encoded through the TRAILBLAZER encoder to obtain their corresponding latent representations. Leveraging the additive structure of the latent space, a per-patient delta vector was computed by subtracting the pre-treatment latent from the post-treatment responder latent, capturing the directional shift associated with a successful treatment response. Cosine similarity was then calculated between these delta vectors and each entry in the perturbation embedding dictionary derived from the mechanism segmentation network, obtaining a ranked list of candidate perturbations most aligned with the observed response trajectory.

### Energy distance plots

The TRAILBLAZER network was trained solely on the cell reconstruction objective, within the PARSE PBMC dataset^30^, using 9 donors for training and 3 held-out donors for validation. Cells are reconstructed by decoding latent representations together with their corresponding sums or subtractions of average perturbation embeddings produced by the mechanism segmentation network. No latent metric losses were applied at this stage. Training was performed on the same PARSE subset described above, with cells grouped into sets of 1, 5, 50, and 500. Across all configurations, exactly 1000 cells are presented to the network in each forward pass, ensuring that every training iteration exposes the model to an equal amount of information regardless of group size. Reconstruction quality was evaluated using Energy Distance (ED), which was chosen over simpler metrics as it compares distributions beyond first moments.

To compute the mean ED during a run, we first sample N random donor-perturbation pairs from the validation split. For each pair, M batches are inferred from the selected donor’s control cells, with the corresponding perturbation embedding summed in latent space so that the network operates as a perturbation inference model. Because the network generates cells in an unordered fashion, output cell types are assigned post-inference using a Support Vector Classifier (SVC) trained on the full PARSE dataset. From each of the resulting output and target cell groups, 25 cells per cell type are randomly sampled. Cell types with fewer than 25 cells are excluded, and only cell types present in both output and target are retained for comparison. The ED is then computed gene-wise across the 25 cells for each cell type, and averaged to yield a mean ED per cell type. A final per-pair summary is obtained by averaging the cell-type-level EDs across all cell types for that donor-perturbation combination.

### Perturbation segmentation analysis

The mechanism segmentation network was trained on PARSE PBMC dataset^30^ to cluster groups of cells by treatment, using context sets of 1, 5, 50, and 500 cells. Across all configurations, batch sizes were chosen such that the network processes an equal number of cell states per forward pass regardless of group size. Network performance is characterized through two complementary geometric measures computed in latent space: average cosine distance, defined as 1 − cosine similarity (where a value of 1 indicates orthogonality between embeddings), and average euclidean distance. Both metrics are reported in two regimes: inter-treatment, measuring distances between groups belonging to different treatments, and intra-treatment, measuring distances among groups belonging to the same treatment, to jointly assess how well the embedding space separates distinct treatments while keeping identical ones together.

To compute the euclidean distances, we first infer cell group embeddings for each treatment and represent each group by its mean embedding, normalized to unit norm. Inter-treatment distances are then obtained by computing all pairwise euclidean distances across treatment groups and averaging. For the intra-treatment case, we restrict the computation to treatments for which at least two group embeddings were sampled during inference, compute pairwise distances within each such treatment, and average across treatments.

Cosine distances follow an analogous procedure. For each treatment, we compute a mean embedding and assemble a full pairwise cosine distance matrix as 1 − cosine similarity. The inter-treatment cosine distance is taken as the average of the off-diagonal entries of this matrix. For the intra-treatment cosine distance, we repeat the same computation restricted to embedding pairs sharing the same treatment label, and again average across treatments.

### UMAPs with and without latent shaping

#### Without latent shaping

After training the TRAILBLAZER network on the task of cell reconstruction, we take 5 random donors and 5 random treatments from PARSE PBMCs dataset^30^. In particular, we take the version of the network trained on sets of 500 cells. We infer batches of each possible pair of donor and treatment through the encoder and obtain the corresponding latents. Finally, we produce a UMAP colored by perturbation label, and another colored by donor label.

#### With latent shaping

We take the previous neural network and continue training it by introducing latent metric losses. In particular we introduce the loss to bring healthy states to the center of the hypersphere (zero norm), along with the cosine similarity loss between the TRAILBLAZER network encoder latents and the perturbation segmentation network embeddings. Then, we infer the same combinations of donor and treatment and produce UMAPs in the same fashion as before, colored by perturbation and donor.

### Zero and few-shot reconstruction benchmarking

To evaluate the generalization capabilities of different architectures to unseen treatments, we designed a held-out benchmark as follows. A single treatment was selected and excluded entirely from the PARSE PBMCs^30^ training set, and each architecture (CellFlow^23^ cfp ver. 0.0.1, STACK^24^ ver. 0.1.2, and LPM^32^ ver. 0.1.0) was trained until the validation loss, computed as the reconstruction error for the held-out treatment, reached a plateau. Inference was then performed to reconstruct all cells from all donors for the excluded treatment.

To assess reconstruction quality, we randomly sampled 10 cells per cell type per donor from both the reconstructed and real datasets, repeating this sampling procedure 10 times per donor. Each set of 10 sampled cells sharing the same cell type and donor was averaged into a single pseudo-bulk profile. Pearson correlation coefficients were then computed between predicted and ground truth pseudo-bulk profiles using the top 100 DEGs for the held-out treatment, and the resulting values were averaged across all cell types to yield a single mean Pearson R per treatment.

To further probe the effect of training data availability, we repeated this procedure while progressively increasing the number of donors included in the training set for the held-out treatment. In these experiments, the remaining donors were held out as the validation set. The full benchmark was repeated for selected treatments (IL-15, IL-10 and IFN-gamma).

### Cell importance analysis

We trained a multicellular binary classifier on scRNA-seq biopsies of patients with non-metastatic, treatment-naive primary invasive carcinoma of the breast before treatment with one dose of pembrolizumab (Keytruda or anti-PD-1)^33,34^, incorporating a self-attention pooling layer to predict responsiveness to anti-PD-1 therapy, defined as TCR clonotype expansion, for samples drawn from naive patients. The self-attention pooling layer assigns a learned weight to each embedded cell prior to pooling, producing a single aggregated embedding used for final classification. Each input sample consisted of 20 cells, balanced across cell types using the specialized dataloader described above, to prevent the introduction of cell type composition biases during training. The per-cell attention weights were subsequently averaged across cell types using 100 samples for each donor, yielding a relative importance score per cell type with respect to the final classification decision.

### Disease ranking

A subset of one million cells was drawn from the CELLxGENE dataset^53,54^, sampled to maximize variability across three metadata axes: tissue type, dataset of origin, and disease state. Using a multicellular classifier analogous to the one described above, we trained the model on disease classification using embeddings produced by each of the candidate foundation model architectures (Geneformer^51^, scGPT^9^, scVI^52^), as provided within the CELLxGENE dataset^53,54^. Model performance was evaluated using the F1 score of patient samples, 25 samples per disease.

### Latent shaping drug rediscovery k-rank

We take random pairs of donors and perturbations from PARSE PBMCs dataset^30^and compute their corresponding latents using the TRAILBLAZER network encoder. Specifically, we pick 10 random perturbations, infer 160 samples (500 cells each) for each unique pair, and calculate the mean latent for the perturbed states. Given a donor and perturbation pair, we then calculate the cosine similarity between the mean latent produced by the network and each of the average treatment embeddings previously obtained from the mechanism segmentation network. Finally, we order the values in decreasing order and the rank is calculated as the index belonging to the correct treatment applied (where ideally, a rank 1 would mean that the perturbation embedding for that treatment had the highest cosine similarity). The ranks are averaged among all picked donor and treatment pairs to get the final metric.

### Information preservation

Using the TRAILBLAZER networks previously trained on PARSE PBMCs^30^ for the task of cell reconstruction on sets of 1, 5, 50 or 500 cells, we proceed to infer the cell set samples for every perturbation present in the dataset. Then, we train the mechanism segmentation network, using sets of 50 cells, on each of the synthetic PARSE datasets, generated by networks trained on 1, 5, 50 or 500 cells, and compare their equal error rate (EER) while training. This metric corresponds to the point at which the false acceptance and false rejection rates are equal, with lower values indicating better discrimination. It was compared across all runs as a measure of how well the segmentation network separates distinct perturbation mechanisms.

## CONTRIBUTIONS

A.G. conceived the project, with input from N.C.; J.N. designed and implemented TRAILBLAZER; J.N. analyzed the PARSE PBMCs and breast cancer datasets with input from A.G., S.S.B., and P.S.; P.S. and J.N. performed benchmarking against CellFlow, LPM and STACK models; J.N. and A.G. wrote the manuscript with contributions from S.S.B. and P.S.; J.N. wrote the methods with contributions from A.G. and P.S.; A.G. supervised the project; All authors read and approved the final manuscript.

## DATA AVAILABILITY

All datasets used in this study are publicly available. The PARSE PBMCs perturbation dataset^30^ is available from its original publication and associated repository. Breast cancer single-cell RNA-seq datasets^33,34^ used for treatment response analyses were obtained from previously published studies and are available through their respective repositories. Human disease single-cell datasets used for phenotypic classification benchmarking were obtained from the Chan Zuckerberg Initiative CELLxGENE Discover data portal^53,54^.

Datasets harmonized using AnuBio’s proprietary style-transfer harmonization pipeline, as well as processed data and model outputs generated during this study are available from the corresponding author upon reasonable request.

## CODE AVAILABILITY

Code used for data processing, model training, and analysis in this study is available at https://anubio.ai/resources accessible under CC BY-NC-SA 4.0 license. The repository contains scripts necessary to reproduce analyses reported in the manuscript, including preprocessing, model training, and evaluation pipelines.

Certain components related to AnuBio’s proprietary style-transfer harmonizer are not publicly released but may be made available to academic researchers upon reasonable request and under appropriate agreements.

